# STING/RANTES Pathway in Airway Epithelium Stimulates Sensitization to *Der p1* in an Asthma Model

**DOI:** 10.1101/2023.07.30.550251

**Authors:** Mayoko Tsuji, Mitsuko Kondo, Akira Nishiyama, Tomohiko Tamura, Ayako Nakamura-Ishizu, Etsuko Tagaya

**Author notes:** **Correspondence:** Mitsuko Kondo M.D.

## Abstract

**Background:** Asthma development can be attributed to various factors, including viral infections. Several pathogen including viruses activate stimulators of interferon genes (STING), and a small amount of STING agonist functions as adjuvants for sensitization to house dust mite (HDM); however, the associated mechanism is unclear. We investigated the role of STING during sensitization to HDM in airway epithelial cells.

**Methods:** Airway epithelial cell STING expression was analyzed using the GEO database. We delivered cyclic-GMP-AMP (cGAMP), a STING agonist to mice intranasally, and sorted epithelial cells and performed RNA-seq. Human airway epithelial cells (HBEpCs) were stimulated using cGAMP *in vitro*. Next, we sensitized mice with cGAMP and HDM, *Der p1* on Day 1, and challenged with HDM on Day 7, and on Day 8, analyzed cytokine/chemokine levels, bronchoalveolar lavage cell fraction, histology, and the number of group 2 innate lymphoid cells (ILC2s) and dendritic cells (DCs). Furthermore, we evaluated the effect of RANTES/CCL5 alone on sensitizing to HDM.

**Results:** Relative to other pattern recognition receptors, *TMEM173*, encoding STING, was highly expressed in HBEpCs, and *RANTES* expression was remarkably upregulated in cGAMP-treated mice. *RANTES*, not *IL-33* or *TSLP*, was also activated by cGAMP in HBEpCs, especially in the presence of H_2_O_2_. Type 2 cytokine/chemokine, eosinophil, and goblet cell metaplasia increased with ILC2 and cDC2 accumulation in cGAMP-adjuvanted HDM-sensitized mice. RANTES alone functioned as an adjuvant for induction of type 2 inflammation in mice.

**Conclusion:** STING was highly expressed in airway epithelial cells. STING/RANTES axis may be a crucial pathway for stimulating asthma sensitization.

## 1 Introduction

Asthma is a common type 2 allergic inflammatory disease, and innate as well as adaptive immune cells, cytokines, and chemokines are involved in its pathophysiology.^1^ There are two phases of asthma onset, sensitization and challenge. Once allergens reach the respiratory tract, dendritic cells (DCs) capture the inhaled allergens and present them to naïve T lymphocytes via major histocompatibility complex (MHC) class II (MHCII) molecules and T cell receptors. In the presence of IL-4, the T lymphocytes differentiate into Th2 lymphocytes, producing IL-4, IL-5, and IL-13, and leading to the development of airway eosinophilia, goblet cell hyperplasia, IgE production, and airway hyperresponsiveness.^2, 3^ Antigens, air pollutants, and microbiomes can stimulate pattern recognition receptors (PRRs) in the airway epithelium to release epithelial cytokines and chemokines. Thereafter, epithelial cytokines, such as IL-33 and TSLP can stimulate group 2 innate lymphoid cells (ILC2s), which then produce IL-5 and IL-13. Additionally, DCs can be stimulated by epithelial cytokines and chemokines, affecting adoptive Th2 immunity.^4–6^

Stimulator of interferon genes (STING) is a 379-amino acid dimetric transmembrane protein that binds to endoplasmic reticulum in the cytosol. It is also one of the PRRs that recognizes pathogen DNA in the cytoplasm. The intrusion of the cytosol at the onset of infection by pathogen DNA it is detected by cyclic GMP-AMP synthase (cGAS).^7^ Upon binding to cytosolic DNA, the active form of cGAS undergoes a conformational change, producing the second messenger cyclic GMP-AMP (cGAMP) from ATP and GTP, which then binds to STING.^8, 9^ Further, once STING is activated by cGAMP, type I interferon is produced. Recently, it was reported that human rhinovirus RNA and coronavirus RNA activate STING via cGAS-like receptors,^10–15^ which constitute one of the sources of sensitizing antigens for asthma.^16–18^ It is also activated by several bacterial pneumonias, such as *Streptococcus pneumonia*.^19^ Therefore, the cGAS-STING axis is a potential target for the treatment of various inflammatory diseases.^20, 21^

Reportedly, 2′3′-cGAMP is an endogenous cyclic dinucleotide-based STING agonist used in mice and humans.^21^ Specifically, STING agonists, which are used as adjuvant for vaccination,^22, 23^ enhance asthmatic inflammation in a house dust mite (HDM)-sensitized model of asthma.^24^ It has also been observed that in mice, STING activation leads to the mobilization of inflammatory cells from the bone marrow to other organs.^25, 26^ However, the relationship between asthma sensitization and STING-activated infection remains unclear.

Regulated upon activation, normal T cell expressed and secreted (RANTES)/CCL5 is a chemokine involved in allergy and infectious diseases, including COVID-19 and influenza, as well as cancers. It mediates eosinophils, lymphocytes, neutrophils, and monocytes chemotaxis.^27–32^ Further, it is expressed in airway epithelial cells from patients with asthma and is capable of recruiting eosinophils via the CCR3 receptor.^33–35^ Additionally, poly (I:C), another pathogenLassociated molecular pattern molecule (PAMP) stimulates RANTES production in airway epithelial cells via TLR3.^36^ Reportedly, STING mutant mice display inflammation and *RANTES/CCL5* overexpression in the lungs.^37^ However, the association between STING and RANTES in asthma has not been fully elucidated. Therefore, in this study, we investigated the role of STING in airway epithelial cells during HDM sensitization.

## Material and Methods

### Human datasets

RNA-seq data for human airway epithelial cell samples from children and adults were downloaded from the Gene Expression Omnibus (GEO) database (https://www.ncbi.nlm.nih.gov/geo/). PAMP coding gene expression was analyzed using GraphPad Prism version 9 (GraphPad Software, San Diego, CA, USA).

### Animal model

Pathogen-free C57BL6n mice weighing 20–30 g were used in all the experiments. The mice were housed in sterile microisolator cages with 12-h light/dark cycles. On Day 1, the mice were intranasally sensitized using 1 μg of 2′3′-cyclic GMP-AMP (cGAMP, 531889; Merck Darmstadt, Germany) and 1 μg of (HDM) (*Der p1*, XBP82D3A2.5; Greer Laboratories, Lenoir, NC, USA). On Day 7, the mice were challenged with 1 μg of HDM. The control mice were administered saline. Then, on Day 8, the mice were sacrificed and tissue samples were collected for analysis.

To elucidate the role of RANTES as an adjuvant during sensitization, the mice were sensitized with recombinant RANTES protein (R&D systems, Minneapolis, MN, USA, 478-MR) and 1 μg of HDM on Day 1. On Day 7, they were challenged with 1 μg of HDM. The next day, they were sacrificed and samples were collected for analysis. The control mice were administered saline. All the experimental protocols were approved by the Ethical Review Committee for Animal Experiments at Tokyo Women’s Medical University (approval number: AE23-059).

### BAL analysis

To collect BAL samples, mice under anesthesia were canulated using a 20-gauge indwelling needle and instilled with 1 ml and 0.8 ml PBS. Thereafter, the BAL cell samples collected were to cytospin-stained, dried on slides, and further stained with May-Giemsa. Images were then captured using a BZ51 fluorescence microscope (Olympus, Tokyo, Japan) and analyzed using the imaging software, cellSens (Olympus).

### Histology

For morphological studies, tissue preparation, immunohistochemistry, lectin staining, and hematoxylin and eosin (H&E) staining were performed as previously described.^38^ Briefly, cryo-sections were incubated with 4% Block Ace (Yukijirushi, Hokkaido, Japan) for 20 min. Thereafter, immunohistochemistry was performed using primary antibodies against Ki67 (clone SP6, rabbit IgG, 1:200; NOVUS, Littleton, CO, USA) and CD45 (clone 30-F11, rat IgG2b kappa, 1:100; BD Biosciences, Franklin Lakes, NJ, USA). For immunofluorescent secondary reagents, Alexa 594-conjugated goat antibodies (R37117; Thermo Fisher Scientific, Waltham, MA, USA) and Alexa 488-conjugated goat antibodies (A11029, Thermo Fisher Scientific) were used. The samples were observed using an LSM710 confocal fluorescence microscope (Zeiss, Heidelberg, Germany). Image analyses were performed using IMARIS (Zeiss) or ImageJ software (National Institutes of Health, Bethesda, MD, USA). Periodic acid-Schiff (PAS)/Alcian blue staining procedures were performed as previously described.^39^

### Flow cytometry

Mice lung samples were extracted, minced, and incubated with 50 μg/ml Liberase TM (05401119001; Merck) and 1 mg/ml DNase (314-08071; Nippon Gene, Tokyo, Japan) in RPMI for 30 min. Thereafter, the digested samples were passed through a 40-μm cell strainer and grinded using the inner cylinder of a syringe. This was followed by hemolysis for 2 min using a lysis buffer (0.8 g NH_4_Cl, 0.2 M NaHCO_3_, and 0.5 M EDTA-2Na), washing, and resuspension in 100 μl of FACS solution containing anti-mouse CD16/32 antibody (93). Finally, the cells were analyzed using the CytoFlex S flow cytometry system (Beckman Coulter, Brea, CA, USA). The following monoclonal antibodies (mAbs) were used for the analysis of DCs: rat mAbs against CD45 (30-F11), F4/80 (BM8), CD11c (HL3), MHCII (M5/114), CD11b (M1/70), and CD103 (2E7). The DC fraction was defined as FSC^high^SSC^high^CD45^+^F4/40^+^CD11c^+^MHCII^+^. Further, the cDC1 fraction was defined as CD11b^-^CD103^+^, while the cDC2 fraction was defined as CD11b^+^CD103^-^.^40^ For ILC2 markers, we used the following antibodies: rat mAbs against CD4, CD8, B220, Gra-1, CD11b, and Mac-1, and against CD45, CD25, Sca-1, and KLRG.^38^ The ILC2 fraction was defined as Lin^-^CD45^+^Sca-1^+^KLRG^+^CD25^+^.^41^ All the mAbs were purchased from BD Biosciences, eBioscience (San Diego, CA, USA), or BioLegend (San Diego, CA, USA).

### Cell culture

Human bronchial epithelial cells (HBEpCs; PromoCell Heidelberg, Germany, C-12640) were cultured in a collagen-coated dish and after 2–3 passages, the cells were pelleted, suspended, and incubated in Pneuma Cult-Ex Plus Medium (Stemcell Technologies, Vancouver, BC, Canada) in a collagen-coated 24-well plate at a cell density of 1×10^5^ cells/cm^2^. After 24 h, the cells were stimulated with 10 μg/ml cGAMP in the presence or absence of 100 μM H_2_O_2_,^42, 43^ followed by incubation with 21% O_2_ and 5% CO_2_ at 37 °C for 4 h, after which the cells were collected and assayed.

### Quantitative RT-PCR (qPCR)

Total RNA was extracted from lung tissues and cultured using an ISOGEN kit (Nippon Gene, Tokyo, Japan). Briefly, first-strand cDNA was generated using ReverTra Ace (Toyobo, Osaka, Japan). Thereafter, qPCR was performed using the Thunderbird SYBR qPCR Next Mix (Toyobo, Osaka, Japan) and the Quant Studio3 Real-Time PCR System (Thermo Fisher Scientific, Tokyo, Japan). PCR was initiated via incubation at 95 °C for 20 s to activate polymerase, followed by 40 cycles of 95 °C for 1 s and 60 °C for 20 s. Gene expression levels were then calculated using the delta-delta Ct method, and normalized based on the expression level of *Gapdh*, which served as the internal control. The target primers used are listed in Supplemental Table 1.

### RNA-seq

Mouse lung epithelial cells were prepared and confirmed as Lin^-^VE-cadherin^-^EpCAM^+^ (Supplemental Figure 1). Thereafter, RNA was isolated as described in the preceding section. To synthesize and amplify cDNA, total RNA (1 ng) was injected into the SMART-Seq v4 Ultra Low Input RNA Kit (Clontech, Palo Alto, CA, USA) for sequencing. RNA-seq libraries were prepared using a Nextera XT kit (Illumina, San Diego, California, USA). Further, single-end 75Lbp sequencing was conducted on theDNBSEQ-G400 (GI Tech) or NextSeq 500 (Illumina) platform.

All subsequent analyses were performed using RaNA-seq (https://ranaseq.eu/index.php).^44^ Specifically, RNA-seq reads were quantified using Salmon software set to single-end default parameters. Differential expression analysis was performed using Wald’s test with a parametric fit in DEqeq2. Gene Set Enrichment Analysis (GSEA) was performed for enrichment and network analyses to reveal the enriched pathways based on the KEGG, REACTOME, WikiPathways, LIPID MAPS, and BIOCYC databases.

### Statistical analysis

All data were presented as mean ± SEM and were analyzed via Mann Whitney’s U test, one-way analysis of variance (ANOVA) with Tukey’s post-test, one-way ANOVA with Kruskal-Wallis test, or two-way ANOVA with Bonferroni’s post-test for significant differences. *p* < 0.05, for the null hypothesis, was considered statistically significant.

## Results

### *TMEM173*, a STING-coding gene, showed the highest expression level among PRRs in human airway epithelium and mice EpCAM^+^ epithelium

STING is highly expressed in the lungs.^45^ However, its expression level compared to those of other PRRs has not yet been determined. In this study, to examine STING expression in the airway epithelium, we analyzed the expression of *TMEM173*. Thus, we observed that *TMEM173* showed the highest expression level among the PRR-coding genes based on GEO databased (Figure 1A) and the analysis of mouse Lin^-^CD45^-^EpCAM^+^ lung epithelial cells (n = 3) (Figure 1B). These findings suggested that STING is the most crucial PRR in the steady state.

**Figure 1.**
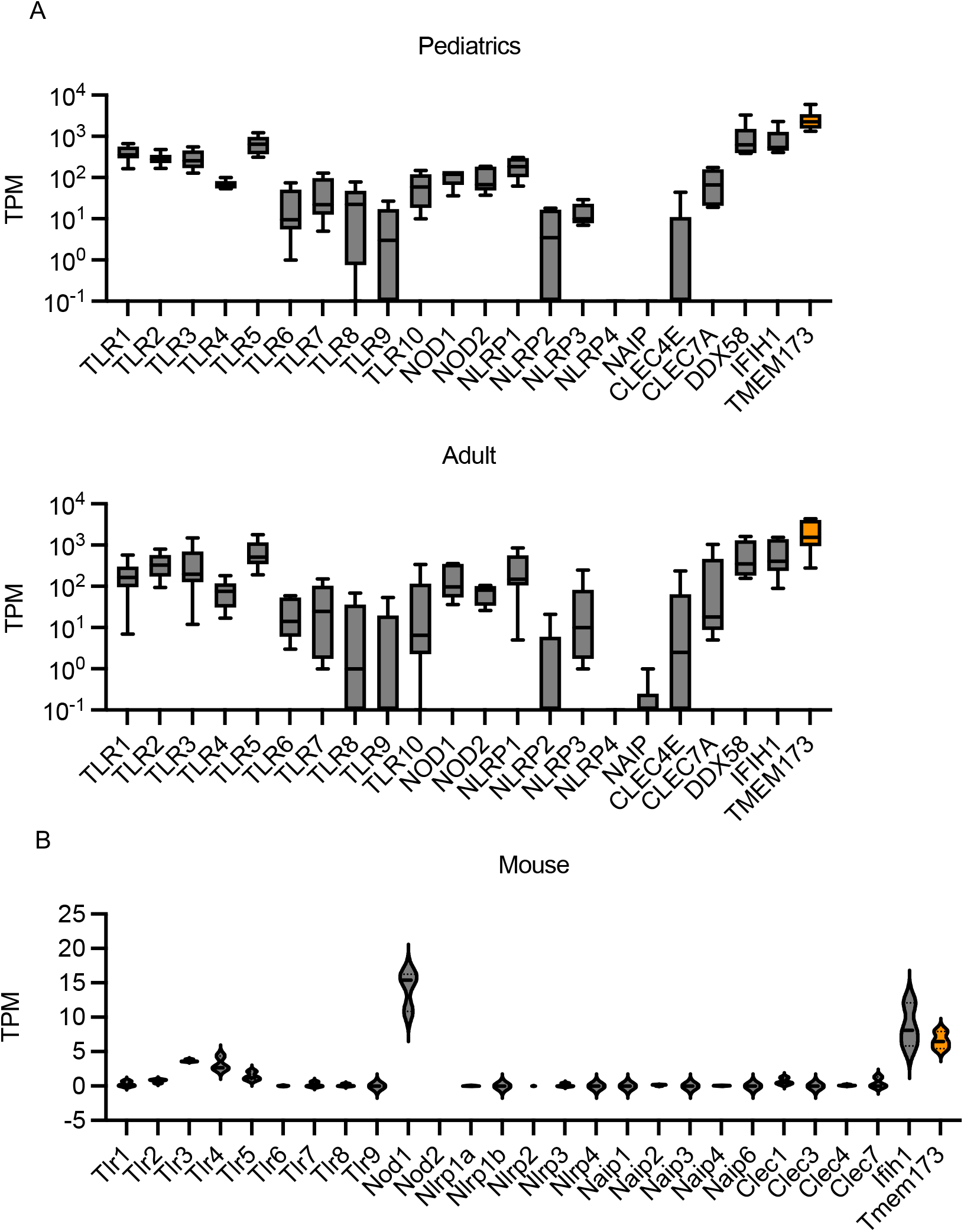
STING expression is the highest of PAMPs in human airway epithelial cells. (A) Expression of pattern-recognition receptor cording genes from pooled RNA-seq data (GSE 148815) of human airway epithelial cells from pediatrics and adults. (B) Expression of pattern-recognition receptor cording genes of Lin^-^VE-cadherin^-^EpCAM^+^ cells in mice.

### STING activation upregulated *RANTES* and accumulated inflammatory cells around bronchi and blood vessels *in vivo*

Mice were intranasally administered cGAMP or PBS once, and BAL, histology, and RNA-seq analyses were performed (Figure 2A). At 12 h following cGAMP administration, we observed increased total cell and neutrophil numbers in BAL in a dose-dependent manner. Lymphocyte numbers tended to increase as well; however, the increase not significant. Further, no significant differences were observed with respect to the number of macrophages or eosinophils (Figure 2B). To observe lung morphological changes, we performed H&E staining on lung tissue sections following the intranasal administrated of cGAMP (50 μg) or PBS. The lung tissue samples exhibited cellular accumulation around the bronchi and blood vessels. To identify the cell type around the blood vessels and bronchus, we performed immunohistochemical analysis on Day 3 on lung tissue sections from mice treated intranasally with 50 μg of cGAMP or PBS. Thus, we observed that the accumulated cells were positive for CD45 (*p* = 0.0079) and Ki67 (*p* = 0.0002) (Figure D), indicating that they were leukocytes proliferating epithelial and endothelial cells.

**Figure 2.**
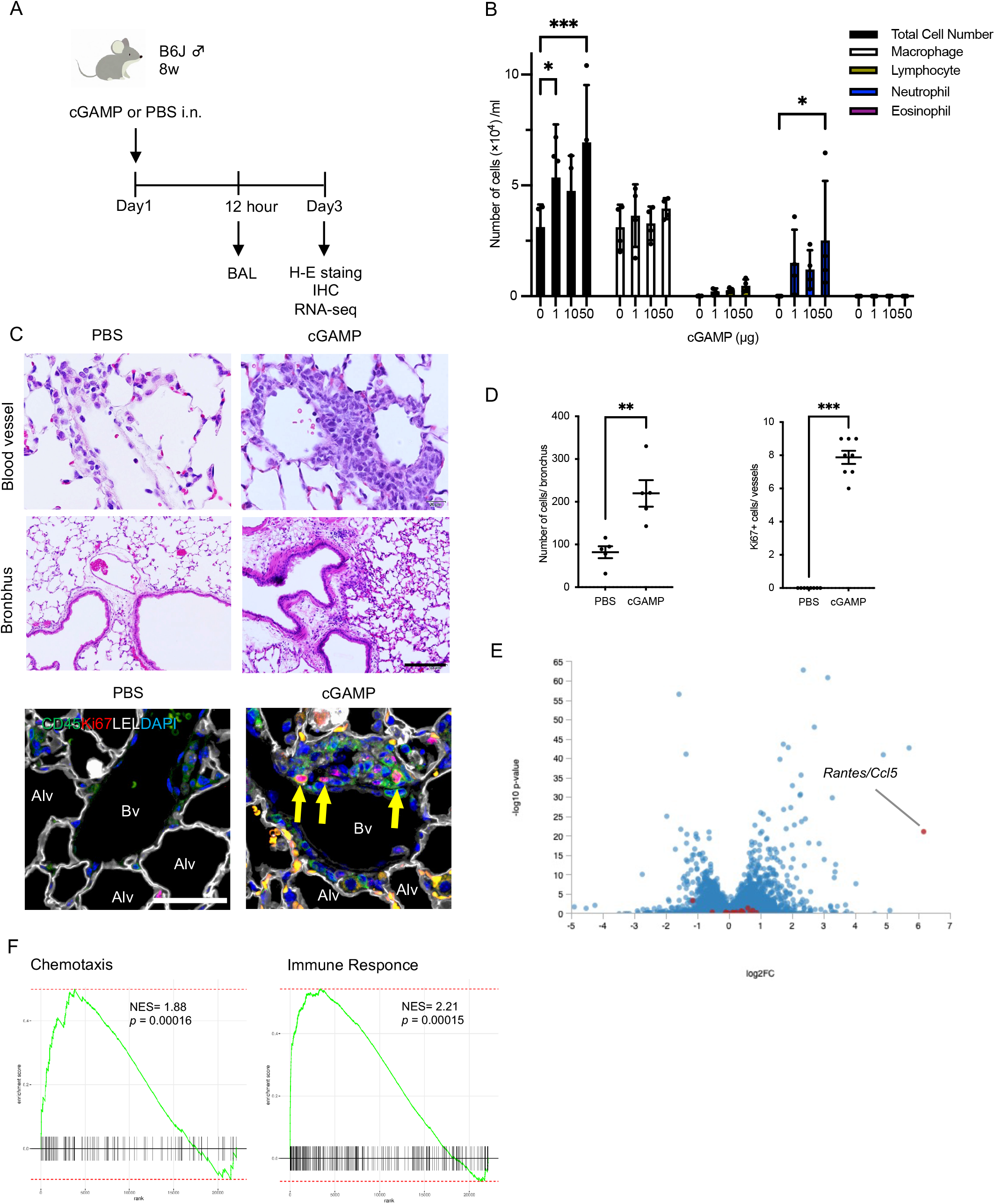
Analysis of STING function with administration of cGAMP *in vivo*. (A) Analysis protocol of stimulating STING by cGAMP. cGAMP was administrated intranasally, then we performed BAL following 12h. Lungs were collected and analyzed on Day 3. (B) RNA-seq analysis of lung CD45^-^VE-cadherin^-^EpCAM^+^ cells 3 days after administrating cGAMP. Ccl chemokines were displayed as red dots. (C) GSEA analysis showing enriched of Chemotaxis and Immune Response related gene sets. (D) Cell fraction in BAL at 12h after administrating 1, 10, and 50 μg or none of cGAMP. Statistical analysis is performed by two-way ANOVA test (means ± SEM, 2 points/ section, n=4). (E) Histological analysis of lung around blood vessels and bronchus after administrating 50 μg of cGAMP, stained with HE. Scale bar: 100 μm. Immunohistochemical analysis of collecting cells in lungs. Green; CD45, Red; Ki67, White; Tomato lectin, LEL, and Blue; DAPI. CD45^+^ cells are accumulated and proliferated around blood vessels (yellow arrow) in cGAMP-administrated lung. Scale bar is 40 μm. (F) Number of cells between bronchus and alveolus analyzed by Image J. Statistical analysis is performed by ordinary Mann-Whitney’s U test (means ± SEM, 2 points/ section, n=3) Numbers of Ki67^+^ cells around vessels in cGAMP-administrated lung. Statistical analysis is performed by ordinary Mann-Whitney’s U test (means ± SEM, 3 points/ n=3).

Next, we stimulated STING in mice via intranasal cGAMP administration, and 2 days after the cGAMP administration, Lin^-^CD45^-^EpCAM^+^ lung epithelial cells were isolated via flow cytometry (Supplemental Figure 2), and RNA-seq was performed. The volcano plot generated from the data obtained showed one of the highest expression levels for *RANTES/CCL5*, especially *Cc* chemokines (red dots) in epithelial cells treated with cGAMP (Figure 2E). The Gene ontology (GO) enrichment analysis based on GSEA revealed the enrichment of genes associated with “Chemotaxis” (GO:0006935, *p* = 0.000158) and “immune response” (GO:0006955, *p* = 0.000145) (Figure 2F).

Further, the enrichment map network obtained based on functional analysis revealed the 25 most significant pathway networks. Particularly, immune system, neutrophil degranulation, innate immune system, antigen processing, and presentation pathways were strongly enriched and showed strong interaction. Furthermore, fatty acid-related pathways were enriched in interactions with the immune pathway (Supplemental Figure 3).

### In airway epithelial cells, *RANTES* was activated by cGAMP and enhanced under H_2_O_2_ condition

In this study, we also analyzed the direct role of STING in airway epithelial cells using 2′3′-cGAMP. Given that STING is localized in the cytosol, some ingenuity is required for its effective binding with cGAMP. Thus, we assessed the role of STING in airway epithelial cells using cGAMP in the presence of H_2_O_2_ *in vitro*.^42, 43^ To this end, HBEpCs were cultured in the presence of cGAMP (14 nM), H_2_O_2_ (100 mM), or cGAMP and H_2_O_2_ for 4 h (Figure 3A). Our results in this regard revealed that under H_2_O_2_ conditions, cGAMP upregulated *interferon be-ta (hIFNb)* expression level relative to the observation made under the control conditions (*p* < 0.0001, Figure 3C); however, the expression level of *INFa* remained unchanged (*p* = 0.64, Figure 3B). Next, we performed qPCR analysis to identify common epithelial cytokines, including *IL-33*, *TSLP*, and *RANTES*. Thus, we noted that *IL-33* (Figure 3D) and *TSLP* (Figure 3E) were not stimulated by cGAMP, whereas *RANTES* (*p* = 0.031, Figure 3F) was significantly stimulated following the cGAMP-only treatment. Furthermore, cGAMP-induced *RANTES* expression was enhanced under H_2_O_2_ conditions (*p* = 0.011, Figure 3F), while those of *IL-33* (Figure 3D) and *TSLP* (Figure 3E) were unaffected. These findings demonstrated that STING activation upregulated *RANTES* in airway epithelium.

**Figure 3.**
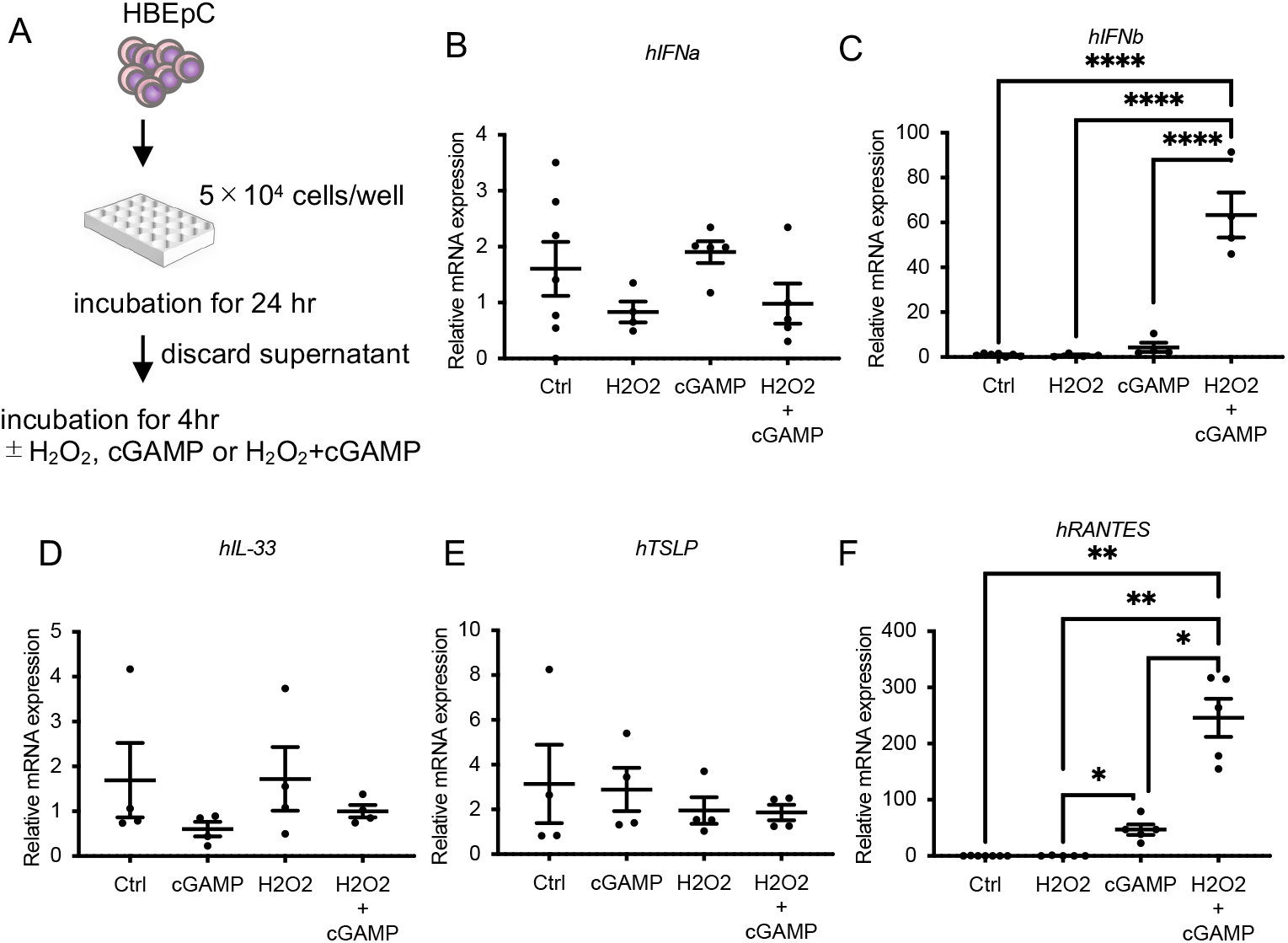
Stimulation of STING induces *RANTES* in human bronchial epithelial cells. (A) HBEpC were cultured with cGAMP (14nM), H_2_O_2_, (100μM) or cGAMP+H_2_O_2_ for 4 hours. (B-F) qPCR analysis of human *IFNα* (B) and *IFNβ* (C) and epithelial cytokines, *IL-33* (D), *TSLP* (E), and *RANTES* (F). *RANTES* was stimulated by cGAMP and cGAMP+ H_2_O_2_. *IFNβ* was also stimulated by cGAMP+H_2_O_2_. Statistical analysis is performed by one-way ANOVA (means ± SEM, n=4). \**p* < 0.05, \*\**p* < 0.01, \*\*\*\**p* < 0.0001

### Administering STING agonist with HDM induced Th2-mediated asthmatic changes

Recently, cGAMP has been used as an adjuvant for vaccination and sensitization to antigens.^22, 23^ Hence, we investigated innate and adaptive immunity and Th2 cytokines in cGAMP-adjuvanted HDM-sensitized mice (cGAMP+HDM). We selected a cGAMP dose of 1 μg, which is the minimal dose required to induce inflammatory changes (Figure 2D). Thus, cGAMP and HDM (1 μg each) were administered intranasally on Day 1, and on Day 7, the mice were challenged with HDM 1 μg. Finally, on Day 8, the mice were sacrificed and samples were collected for analysis (Figure 4A). cGAMP+HDM mice histologically displayed cell infiltration around blood vessels and bronchi, and goblet cell hyperplasia was observed in H&E-stained specimens (Figure 4B). Goblet cells also produced large amounts of mucus in the lung specimens stained with PAS/Alcian blue (Figure 4C) and the BAL cell fraction showed an increase in total cells and eosinophils in the cGAMP-adjuvanted HDM-sensitized mice (Figure 4D). Moreover, qPCR analysis of Th2 cytokines showed elevated *Il-4* (*p* < 0.0001), *Il-5* (*p* = 0.048), and *Il-13* (*p* < 0.0001) levels (Figure 4E). Conversely, HDM-sensitized mice in the absence of cGAMP showed no significant inflammatory changes in BAL and lung specimens.

**Figure 4.**
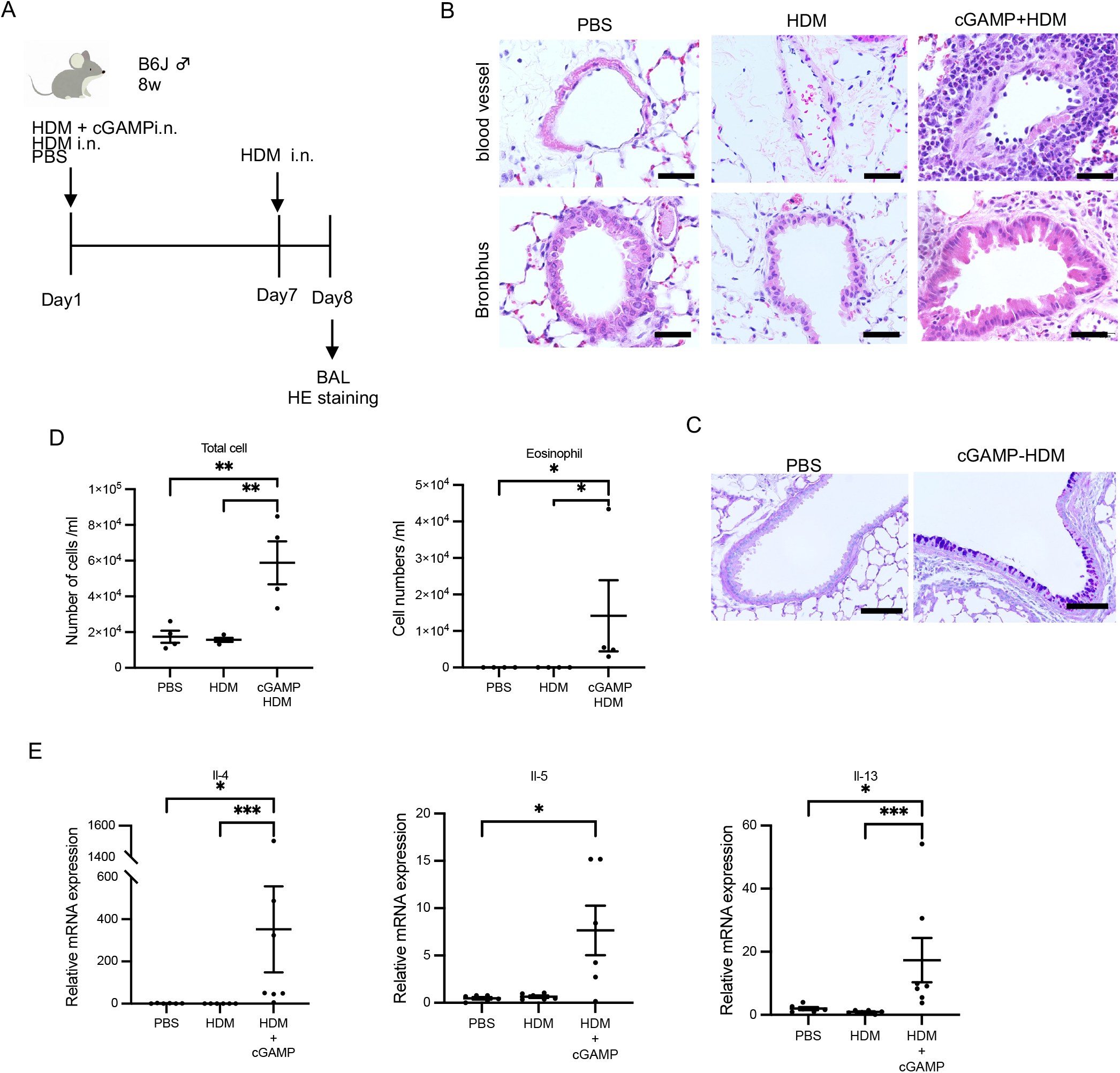
Small amount of cGAMP functions as adjuvant in HDM-sensitized asthma model. (A) Method for HDM-sensitized asthma model. 8-week mice were treated with 1μg of cGAMP and 1μg of HDM, 1μg of HDM or PBS intranasally on Day 1. Mice were challenged with 1μg of HDM intranasally on Day 7, then lungs were extracted and analyzed on Day 8. (B) Pictures of lung sections of control and cGAMP-adjuvanted, HDM-sensitized mice stained with HE. Cells were gathered around pulmonary vessels in cGAMP-adjuvanted, HDM-sensitized mice lung compared to control. Cells were also gathered around bronchus in cGAMP-adjuvanted, HDM-sensitized mice compared to control. Scale bars: 40 μm. (C) Pictures of lung sections in PBS and cGAMP-adjuvanted, HDM-sensitized mouse stained with PAS/Alcian blue. Scale bar: 100 μm. (D) Numbers of Total cell and eosinophils in BAL in PBS-, HDM- and cGAMP-adjuvanted, HDM-sensitized mice. n=4, \*\*\**p* < 0.001. (E) qPCR analysis of lungs in PBS-, HDM -, or cGAMP-adjuvanted, HDM-sensitized (cGAMP+HDM) mice. *Il-4*(D), *Il-5* (E)*, and Il-13* (F). Statistical analysis is performed by one-way ANOVA with Kruskal-Wallis test. \**p* < 0.05, ** *p* < 0.01, *** *p* < 0.001.

### ILC2 and cDC2 proliferated in cGAMP-adjuvanted HDM-sensitized mice

To clarify the connection between innate immune response and adoptive response, we analyzed ILC2s from the lungs using FACS. Our results indicated an increase in the levels of ILC2s in cGAMP-adjuvanted HDM-sensitized mice compared to their levels in PBS-treated HDM-sensitized mice (Figure 5A and B, *p* = 0.0286). This observation suggested that adaptive Th2 asthmatic responses upregulate ILC2s.

**Figure 5.**
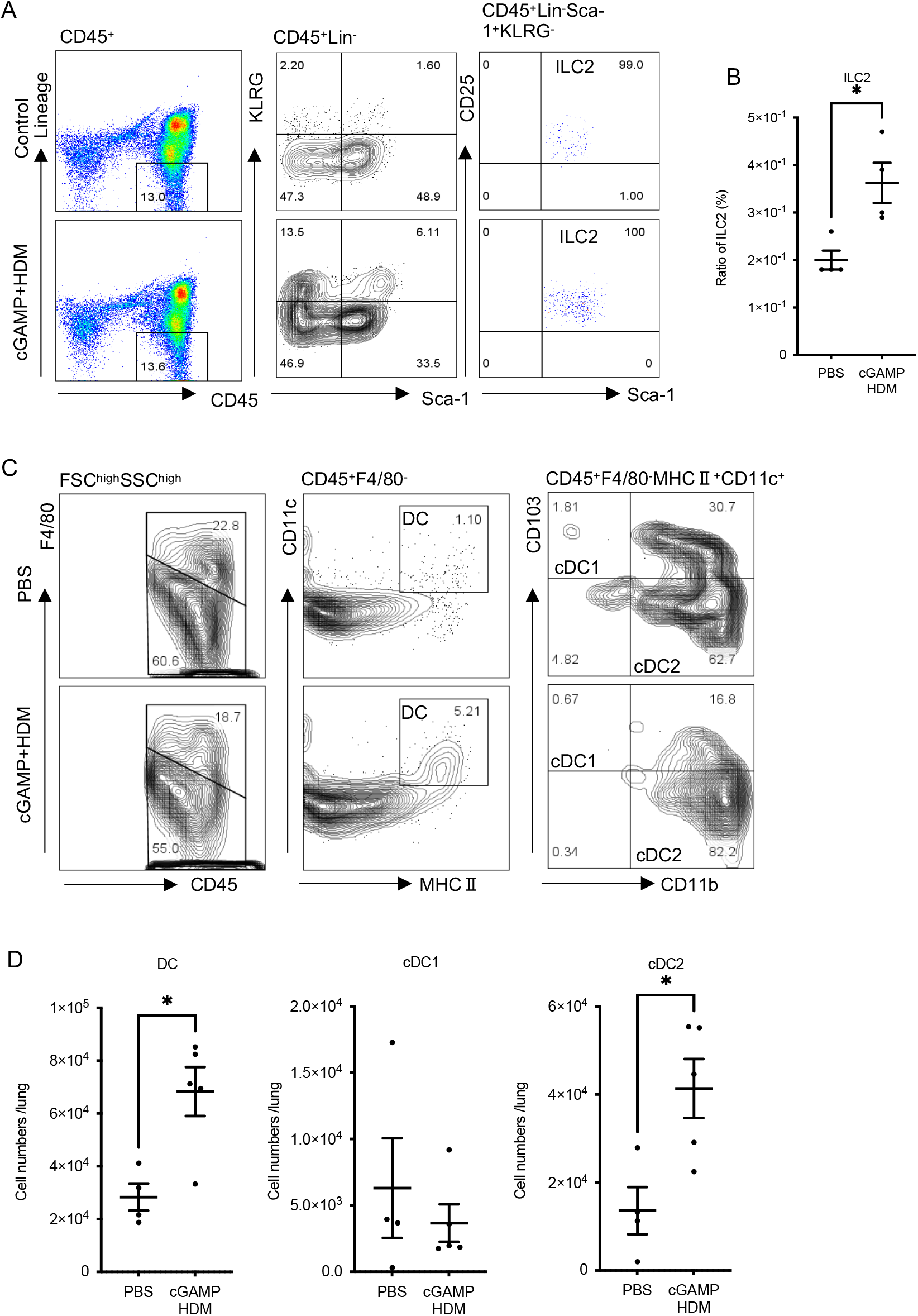
Type 2 inflammation is accelerated in cGAMP-adjuvanted, HDM-sensitized mice. (A) Gating strategy of ILC2s. ILC2 fraction is defined as CD45^+^Lin^-^Sca-1^+^KLRG^-^CD25^+^ fraction. (B) Numbers of ILC2 in lung of cGAMP-adjuvanted, HDM-sensitized mice compared to control. Statistical analysis is performed by ordinary Mann Whitney’s U test (means ± SEM, control; n=4) (C) Gating strategy of DCs. DC fraction is defined as FSC^high^SSC^high^CD45^+^F4/40^+^CD11c^+^MHCII^+^. cDC1is defined as CD11b^-^CD103^+^ DC fraction. cDC2 is defined as CD11b^+^CD103^-^ fraction. (D) Numbers of DC, cDC1, cDC2 in lung of cGAMP-adjuvanted, HDM-sensitized mice compared to control. Statistical analysis is performed by ordinary Mann Whitney’s U test (means ± SEM, control; n=5, +anti-CD103 Ab; n = 6)

Conventional type2 DCs (cDC2s) constitute a DC fraction that recognizes allergens and primarily accelerates Th17 differentiation and Th2 inflammation, whereas cDC1s are less involved in Th2 inflammation. Thus, we analyzed these two types of cDCs. The total DC and cDC2 numbers were increased in cGAMP-adjuvanted HDM-sensitized mouse lungs, whereas the number of cDC1 remained unchanged (Figure 5C and D). These findings demonstrated that cDC2, but not cDC1, is activated in cGAMP-adjuvanted HDM-sensitized mice.

### RANTES functions as an adjuvant in HDM-sensitized mice

In this study, we investigated RANTES expression in STING-activated airway epithelial cells (Figure 3). Reportedly, RANTES mediates the chemotaxis of eosinophils, lymphocytes, neutrophils, and monocytes and induces asthma.^27–31^ To investigate RANTES function during sensitization, we administered 0, 10, and 200 ng of RANTES to mice intranasally and thereafter, determined the effective dose required to evoke inflammatory cells in BAL fluid (Supplemental Figure 4). Given that 10 ng of RANTES significantly induced inflammation, we administrated 1 μg of HDM+PBS or 1 μg of HDM+10 ng of RANTES intranasally, and on Day 7, challenged the mice with HDM (Figure 6A). Thus, we observed inflammatory cell infiltration around the bronchi and blood vessels in the lungs of HDM + RANTES mice, in contrast to the lungs of HDM + PBS mice (Figure 6B and C). Most of the cells in the samples from HDM + RANTES mice were eosinophils. To check for goblet cell hyperplasia, we performed PAS/Alcian blue staining, which revealed goblet cell hyperplasia in the lungs of HDM + RANTES mice (Figures 6D and E). These findings suggested that RANTES acts as an adjuvant for HDM sensitization.

**Figure 6.**
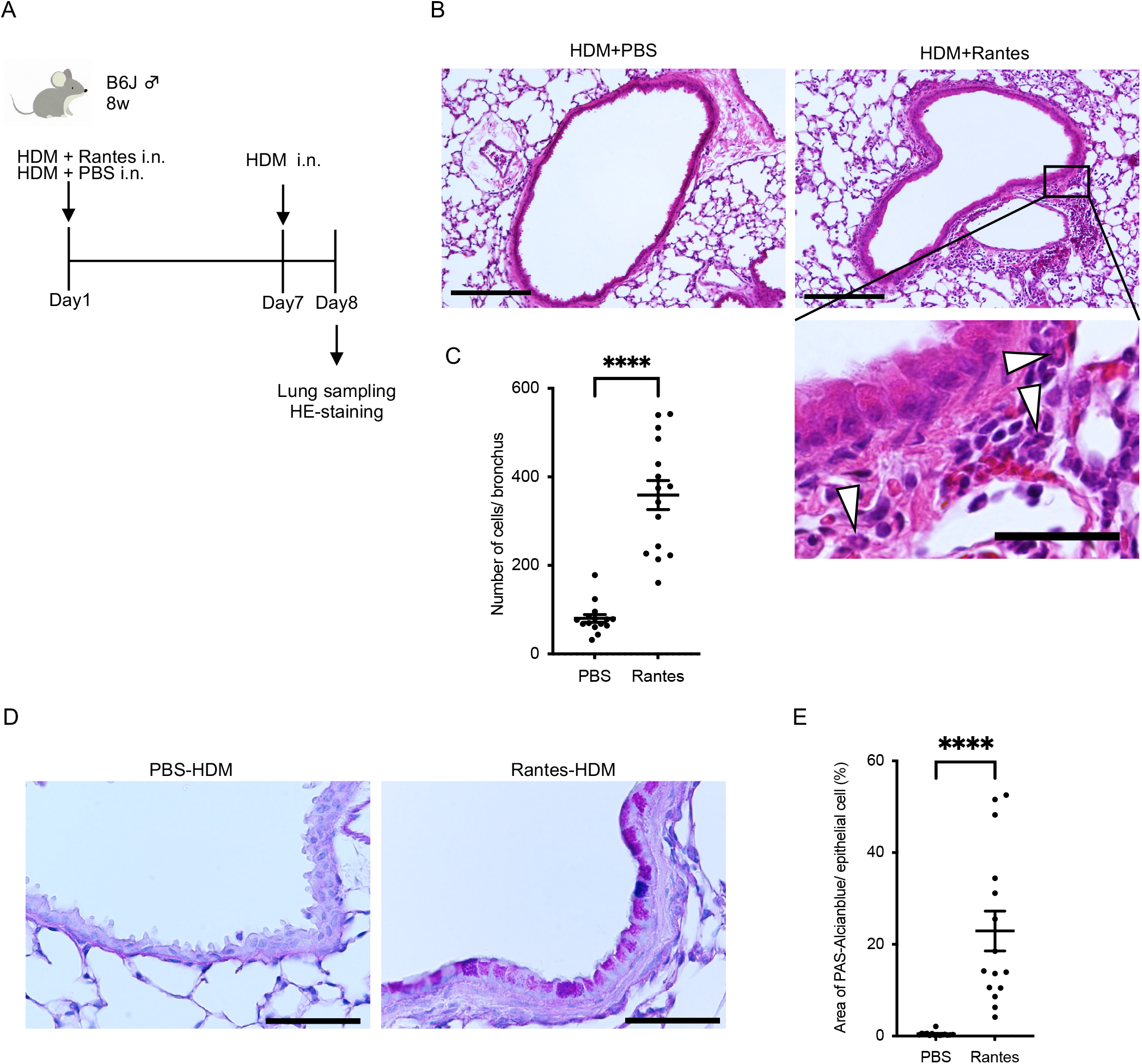
RANTES is the principal role as an adjuvant in sensitization of STING-related asthma development. (A) 8 weeks mice were treated with 20 ng of RANTES with 1 μg of HDM or 1 μg of HDM intra nasally on Day 1. Mice were challenged with 1 μg of HDM intranasally on Day 7, then lungs were extracted and analyzed on Day 8. (B) Pictures of lung sections of control and RANTES-adjuvanted, HDM-sensitized mice stained with H-E. Scale bar: 200 μm. Scale bar in picture of high magnification view: 40 μm. (C) Number of cells between bronchus and alveoli analyzed by ImageJ. Statistical analysis is performed by ordinary Mann Whitney’s U test (means ± SEM, 3 points/ section, n=5). (D) Pictures of lung sections in PBS and RANES-adjuvanted, HDM-sensitized mouse stained with PAS/Alcian blue. Scale bar: 80 μm. (E) Ratio of PAS/Alcianble area per epithelial cells (%). **** *p* < 0.0001

## Discussion

This study showed that among the PRRs in human airway epithelial cells from both children and adults, *TMEM173*, a STING coding gene, showed the highest expression level, and that airway epithelial RANTES produced by STING stimulation may be the core mechanism of sensitization to HDM. These findings suggest that the STING/RANTES pathway plays an important role in the development of HDM sensitization in asthma.

Pathogenic DNA penetrates the cytoplasm and binds to cGAS, producing cGAMP, and RNA also activates STING via cGAS like receptor.^14, 15^ Additionally, cGAMP is transported by SLC19A1 to STING, which activates the interferon cascade and produces type I interferons, such as IFNb.^46^

The production of IL-25, IL-33, and TSLP by airway epithelial cells following injury owing to factors, including air pollution, tobacco, and infection, leads to the activation of ILC2s, which produce Th2 cytokines and induce type 2 inflammation.^3^ Our RNA-seq data showed that *RANTES/CCL5* was highly upregulated in cGAMP-treated lung epithelial cells. We also found that cGAMP administration directly induced *RANTES* expression in airway epithelial cells, while common epithelial cytokines, such as *IL-25*, *IL-33*, and *TSLP*, did not show such response. *In vitro*, *RANTES* also showed enhanced expression in the presence of H_2_O_2,_ an airway epithelial injury factor.^42, 43^ Given that STING exists in the cytosol, its binding to cGAMP may be accelerated by H_2_O_2_, which causes airway epithelial cell damage. Similarly, *IFNb* expression in airway epithelial cells was upregulated by cGAMP treatment in the presence of H_2_O_2_ (Figure 3C). This finding supported the hypothesis that STING activation occurs under H_2_O_2_ in this system. H_2_O_2_ is a reactive oxygen species produced by neutrophils in a broad range of lung diseases, including asthma and viral infection.^47, 48^ Our *in vivo* findings based on BAL analysis and the results of experiments with cGAMP or RANTES alone confirmed neutrophilic inflammation. Additionally, pathway analysis showed that neutrophil degranulation was strongly enriched in cGAMP-treated mice, suggesting that cGAMP-induced RANTES production was enhanced by neutrophil accumulation. Taken together, we speculated that cGAMP stimulated RANTES production in airway epithelial cells, resulting in leukocyte accumulation around the airway and blood vessels, and further RANTES production owing to H_2_O_2_, was possibly led to the release of leukocytes.

Several studies have demonstrated that the cGAS/STING pathway activates DCs, macrophages, neutrophils, and T cells.^11, 19, 49^ GSEA revealed that chemotaxis- and immune response-related genes were enriched in cGAMP-treated epithelial cells (Figure 2C). In contrast, RANTES typically collects leukocytes, such as DCs, neutrophils, lymphocytes, macrophages, and eosinophils.^29–31, 33–35^ Additionally, our results showed lymphocyte proliferation in BAL following cGAMP administration (Supplemental figure 5). Therefore, it is possible that the cells that accumulated around the bronchi and blood vessels in the cGAMP-treated lungs did not only contain neutrophils, but also macrophages, DCs, and lymphocytes. Therefore, further investigations are required for clarification in this regard.

DCs are classified under two subsets, plasmacytoid DCs (pDCs) and cDCs, and based on their residency and roles, cDCs are further classified as cDC1 and cDC2. cDC1 is positive for CD103, while cDC2 is positive for CD11b. Additionally, CD103 is an integrin and its ligand is E-cadherin, which is specifically expressed in the epithelium.^50–54^ Our results showed that ILC2 and cDC2 proliferated in the lungs of cGAMP-adjuvanted HDM-sensitized mice (Figure 5). It has also been reported that these cells are associated with type 2 inflammation.^55^ Although RANTES has been reported to mobilize cDC1 in cancer immunology,^56–58^ we observed the accumulation of cDC2 in our asthma model. These observations indicated that RANTES plays a role in the accumulation of disease-specific leukocytes. Recently, it was reported that cDC2 induces Th2/ Th17 differentiation in an asthma model,^59^ while the deletion of CD103^+^DCs worsens airway allergic response.^53^ The STING/RANTES pathway enhances adoptive immunity via cDC2 accumulation, as well as innate immunity via ILC2 accumulation, suggesting its readiness for future allergic exacerbation.^60–62^ Via RNA-seq, we found that antigen processing and presentation signaling were highly enriched in cGAMP-treated mice using. Taken together, the STING/RANTES pathway may play a key role in the expression of adjuvants during infection-induced sensitization. The limitation of this study is that it has not demonstrated whether RANTES alone is essential or partially involved in STING activated allergen sensitization. Further investigation is needed. Interestingly, the recent paper has reported that RANTES/CCL5 links type1 neutrophilic inflammation to type 2 inflammation in an asthma subgroup ^63^ This supports the importance of RANTES in the bridge between STING and asthma.

In conclusion, our results demonstrated that airway STING activation led to leukocyte accumulation via RANTES production, which increased under H_2_O_2_-induced injury *in vitro*. These findings suggested that RANTES is a key player in the adjuvant mechanism intiating sensitization to HDM and that STING, the most highly expressed PRR, is often involved in the onset of asthma triggered by airway infection. Thus, RANTES may be a new therapeutic target for the development of asthma resulting from STING-activated infections.

## Acknowledgements

The authors thank Masayuki Shino, Yoshimi Sugimura, and Atsuko Ohnishi for their technical support.

This work was supported by JSPS KAKENHI Grant Number JP20K17198, in part by a research grant from TWMU Career Development Center for Medical Professionals, and The Japanese Respiratory Society Research Grants Program A. And this research was supported by MEXT Promotion of Distinctive Joint Research Center Program Grant Number JPMXP0622717006 at the Advanced Medical Research Center, Yokohama City University.

This study was supported by Institute for Comprehensive Medical Sciences (ICMS) and Institute of Laboratory Animals (ILA), Tokyo Women’s Medical University.

## Notes

### Competing Interest Statement

The authors have declared no competing interest.

