## Supplemental figures for "STING/RANTES Pathway in Airway Epithelium Stimulates Sensitization to *Der p1* in an Asthma Model"

| Gene | Forward primer (5'→3') | Reverse primer (5'→3') |
| --- | --- | --- |
| **Human** | | |
| *hGAPDH* | TGTGGGCATCAATGGATTTGG | ACACCATGTATTCCGGGTCAAT |
| *hIL25* | CAGGTGGTTGCATTCTTGGC | GAGCCGGTTCAAGTCTCTGT |
| *hIL33* | GTGACGGTGTTGATGGTAAGAT | AGCTCCACAGAGTGTTCCTTG |
| *hTSLP* | ATGTTCGCCATGAAAACTAAGGC | GCGACGCCACAATCCTTGTA |
| *hRANTES* | GCATCTGCCTCCCCATATTC | CAGTGGGCGGGCAATG |
| *hIFNa* | GCCTCGCCCTTTGCTTTACT | CTGTGGGTCTCAGGGAGATCA |
| *hIFNb* | ATGACCAACAAGTGTCTCCTCC | GGAATCCAAGCAAGTTGTAGCTC |
| **Mouse** | | |
| *mGapdh* | AGGTCGGTGTGAACGGATTTG | TGTAGACCATGTAGTTGAGGTCA |
| *mIl13* | TGAGCAACATCACACAAGACC | GGCCTTGCGGTTACAGAGG |
| *mIl33* | GGCCTTGCGGTTACAGAGG | AACGGAGTCTCATGCAGTAGA |
| *mTslp* | TTCACTCCCCGACAAAACATTT | TGGAGATTGCATGAAGGAATACC |
| *mEotaxin* | GAATCACCAACAACAGATGCAC | ATCCTGGACCCACTTCTTCTT |
| *mRantes* | GCTGCTTTGCCTACCTCTCC | TCGAGTGACAAACACGACTGC |

**Supplemental table 1. Primer sequence for qPCR**

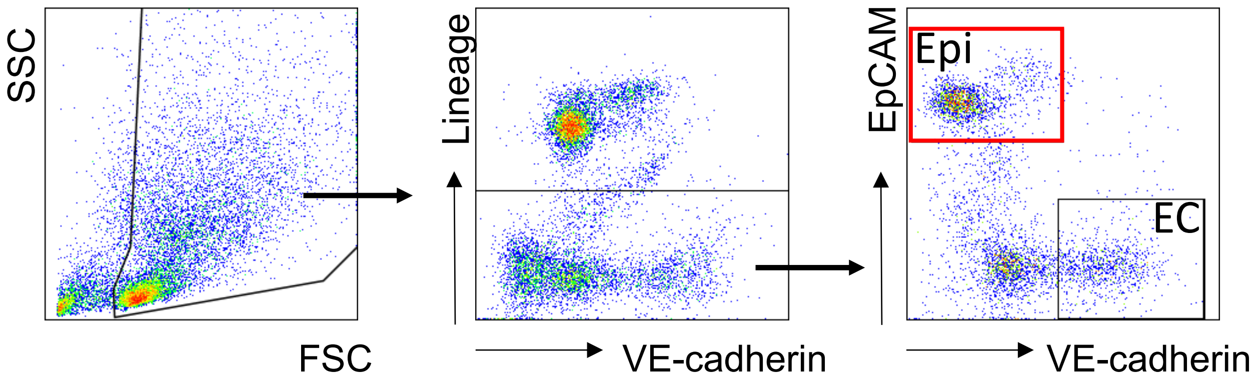


**Supplemental figure 1. Lung Epithelial cell sort gate for RNA-seq.**

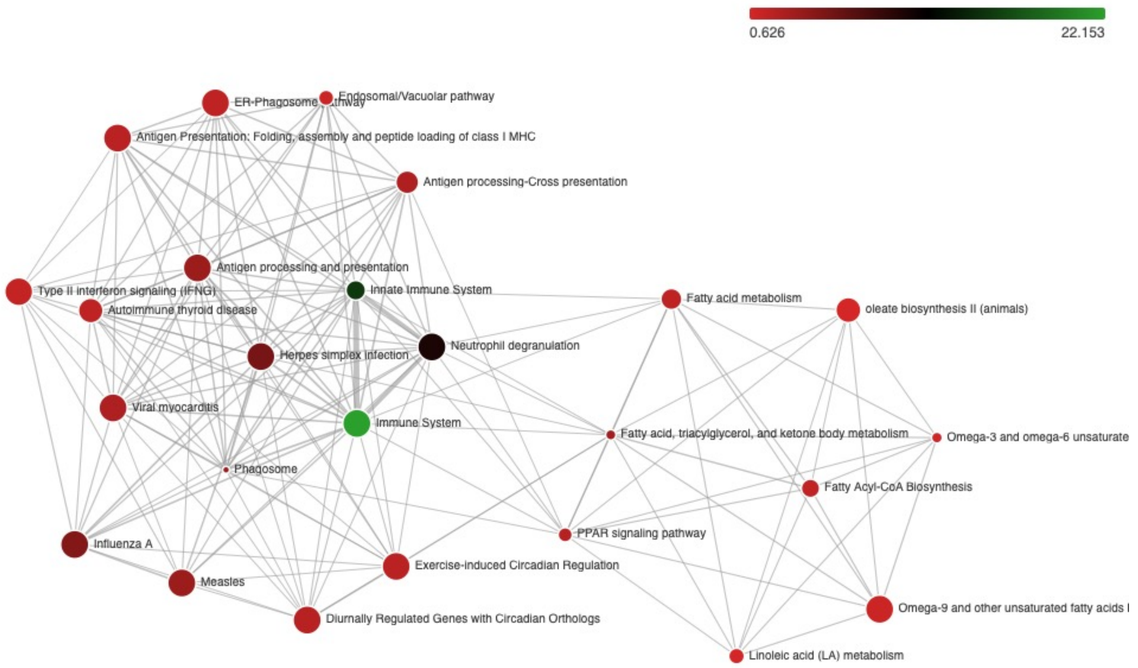


**Supplemental figure 2.** Enrichment map network of functional analysis of $Lin^{-}$VE-$cadherin^{-}$ $EpCAM^{+}$ lung cells in cGAMP administrated mice.

Enrichment map network of functional analysis resulting in the top 25 significant pathway network.

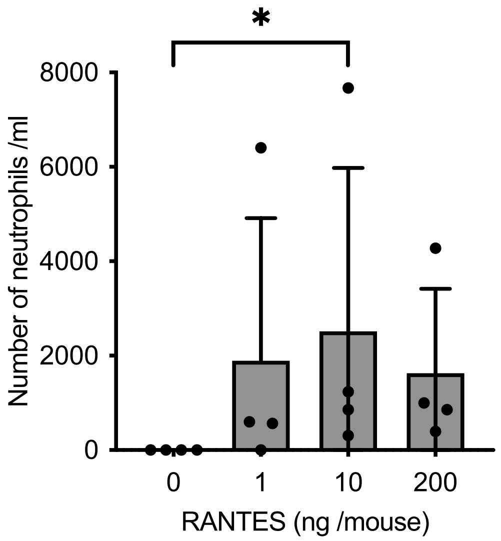


**Supplemental figure 3 Numbers of neutrophil in BAL after administrating RANTES.**

To determine the dose of RANTES sensitization, we administrated 1, 10, and 200 ng / mice of RANTES intranasally. After 20 hours, we performed BAL. we counted the number of neutrophils. (n=4) Statistical analysis was performed by ordinary one-way ANOVA test.

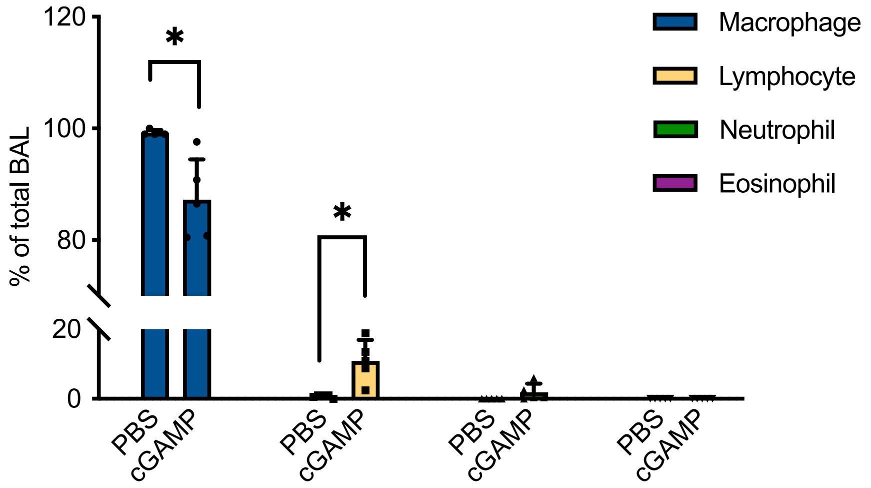


**Supplemental figure 4. The ratio of lymphocytes in BAL were increased 3 days after administrating cGAMP.**

To determine the dose of RANTES sensitization, we administrated 50 µg of cGAMP intranasally. After 3 days, we performed BAL. We counted the number of macrophages, lymphocytes, neutrophils, and eosinophils (n=5) Statistical analysis was performed by ordinary two-way ANOVA test.
